## Supplemental figures for "Long-term tracking of social structure in groups of rats"

### Supplementary information:
### Long-term tracking of social structure in groups of rats

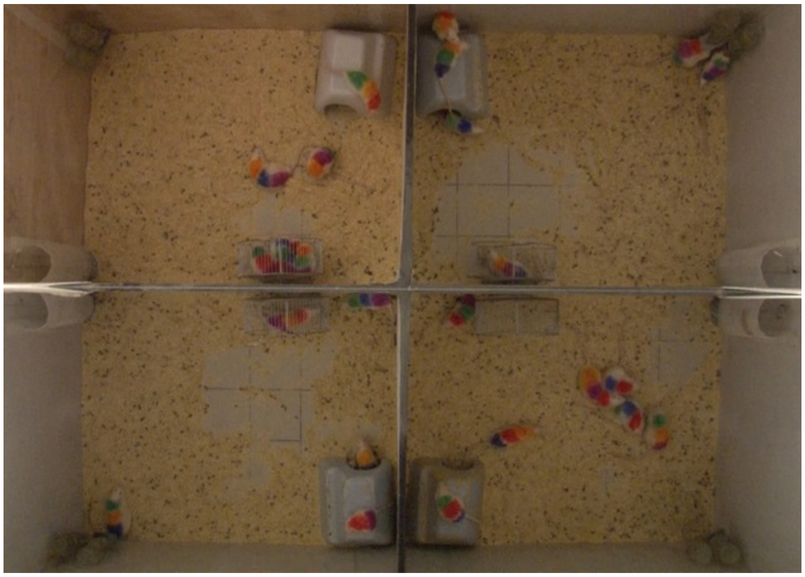

**Figure S1.** **Example camera frame image**. Taken at low-light condition (i.e. during active period).

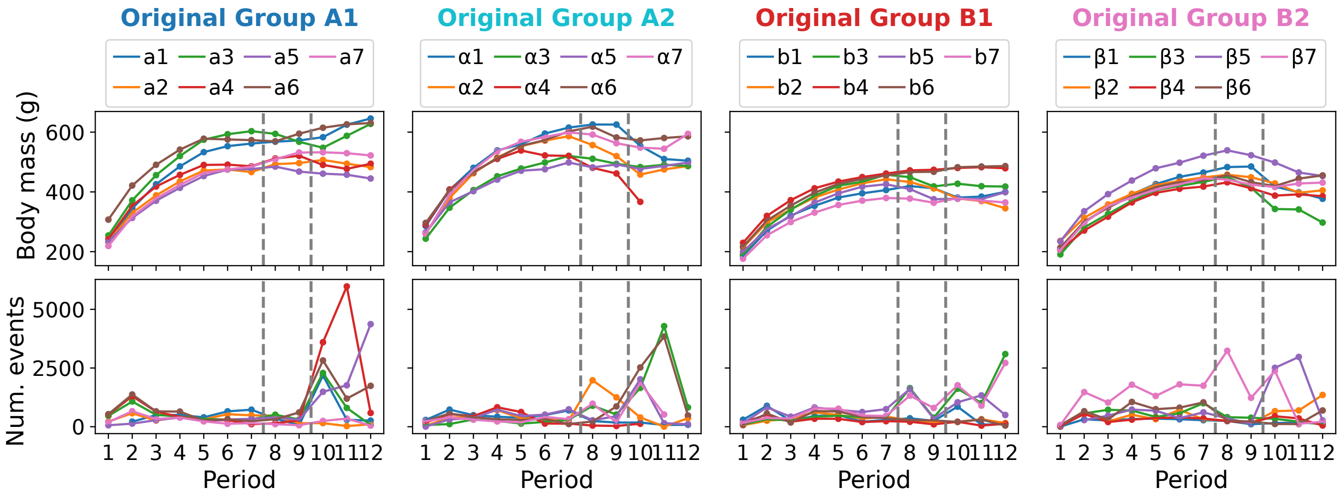


**Figure S2.** **Body mass and number of events during the entire experiment**. Individual rats are plotted according to phase 1 groups and shown with different colored lines. Num. events are the mean number of pairwise events one rat had with other rats in the same group.

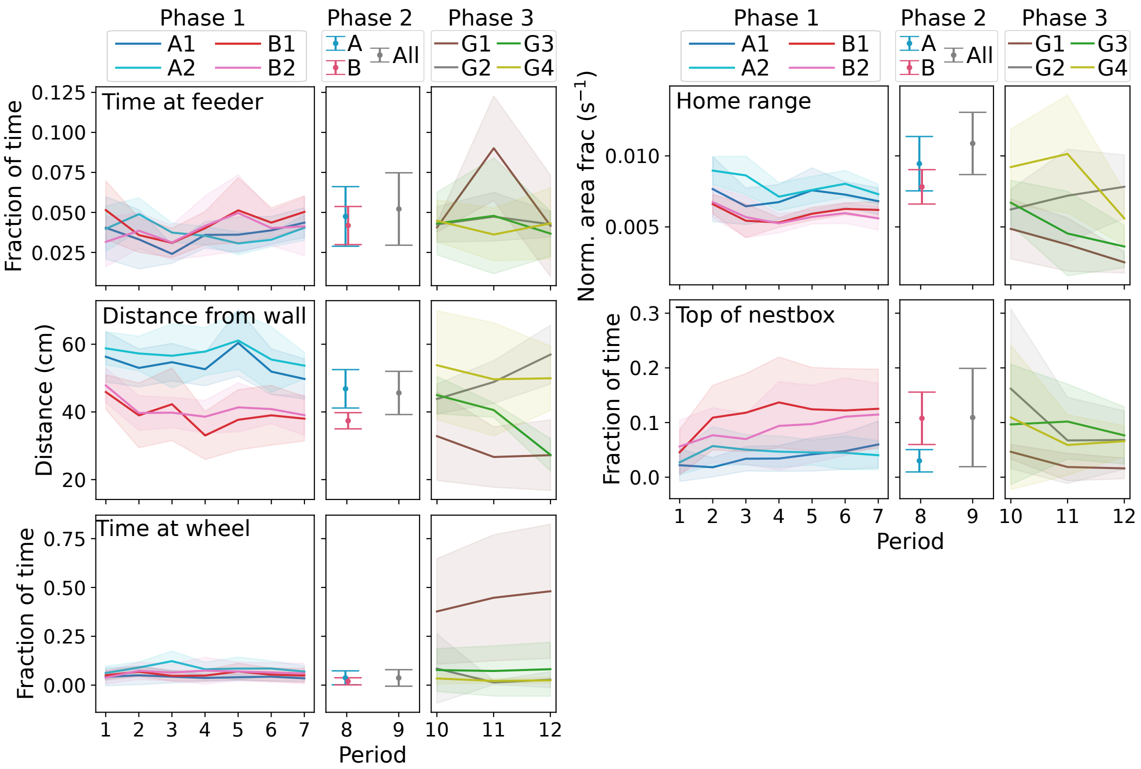


**Figure S3.** **Space use metrics for each group**. Metrics of time at feeder, distance from wall, home range, top of nestbox, and time at wheel. The lines/shaded area or points/error bars show the mean/standard deviation of each metric within each group, for the different phases. Note: home range was not calculated for Pd 1.

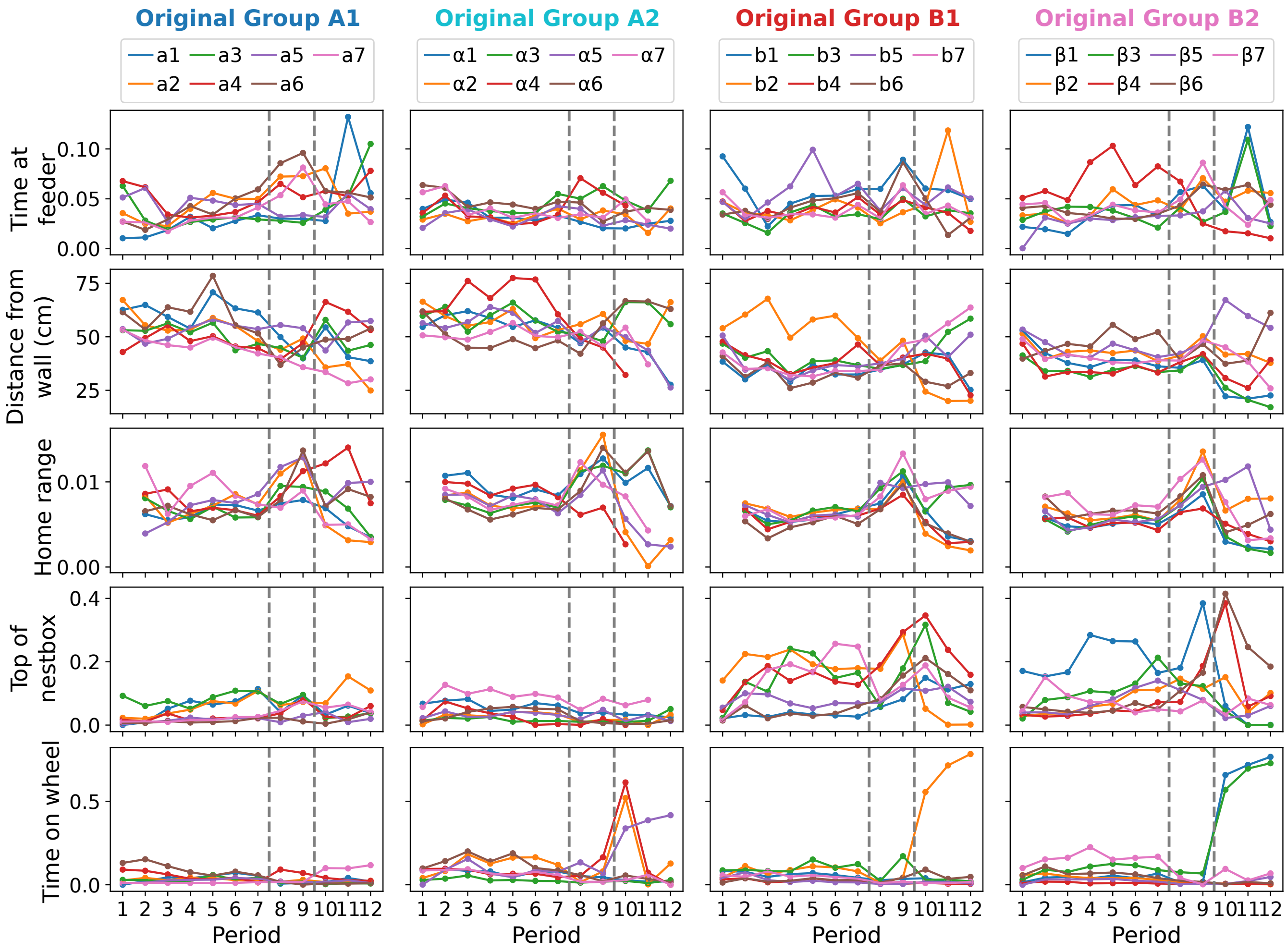


**Figure S4.** **Space use metrics plotted for individual rats**. See also Fig S3 for the average and standard deviation of space use metrics across individual rats in each group. Rats are plotted according to phase 1 groups in an analogous way to Fig S2. Time at feeder is calculated as fraction of time, distance from wall has units of cm, home range is calculated as normalized area fraction ($s^{-1}$), and top of nestbox and time on wheel are are calculated as fraction of time.

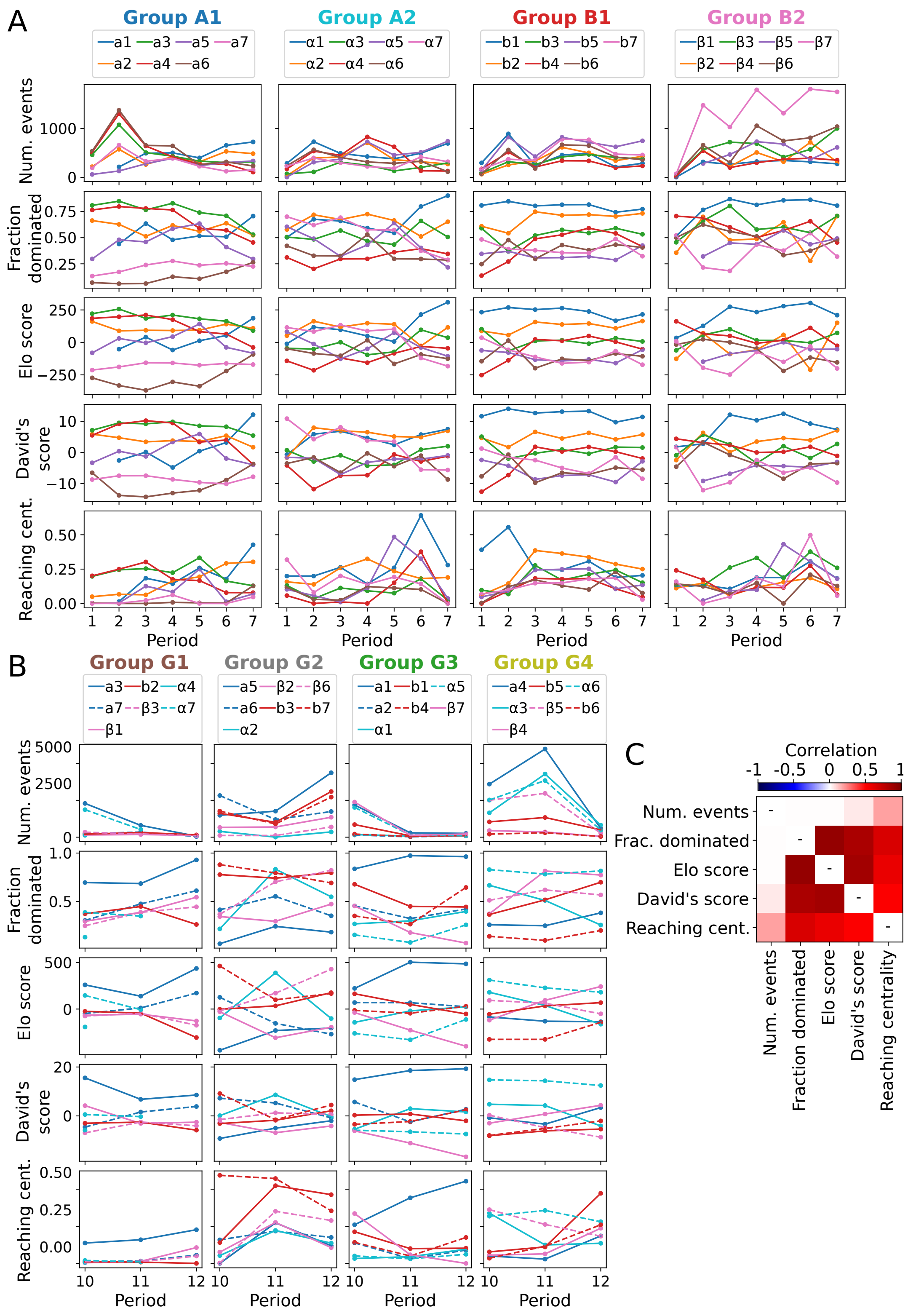


**Figure S5. Individual ranking metrics comparison and correlations.** Shown are the total number of events for each rat, and metrics based on the matrix of pairwise event outcomes for rats in each group, including fraction of events dominated, Elo score, David's score, and the Local Reaching Centrality (LRC). (A) Values for each rat according to phase 1 groups. (B) Values for each rat according to phase 3 groups. Colors designate which groups each rat belonged to during phase 1. (C) Correlation among metrics, calculated as the correlation coefficient for a given pair of metrics during the same period, using all data from phases 1 and 3.

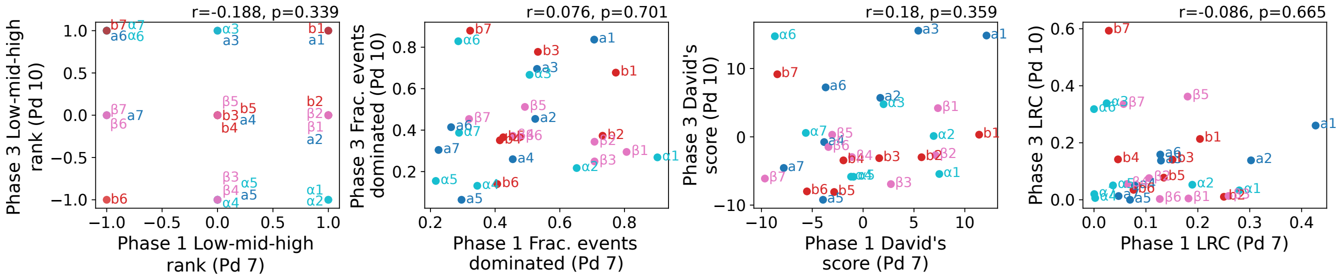


**Figure S6.** **Other phase 1 to phase 3 individual social ranking measures.** See Fig 6 for phase 1 to phase 3 Elo scores for individual rats. Shown here are ranks based on Elo scores as well as other individual rat ranking metrics, including fraction of events dominated, David's score, and local reaching centrality (LRC). Each plot compares scores at the end of phase 1 (Pd 7) to those at the start of phase 3 (Pd 10), i.e. after the new groups were formed. Low-mid-high ranks are determined by assigning a subordinate (low) value of -1 to the lowest two Elo scores in a group during a certain period, a dominant (high) value to the highest two Elo scores, and a middle (0) value to others.

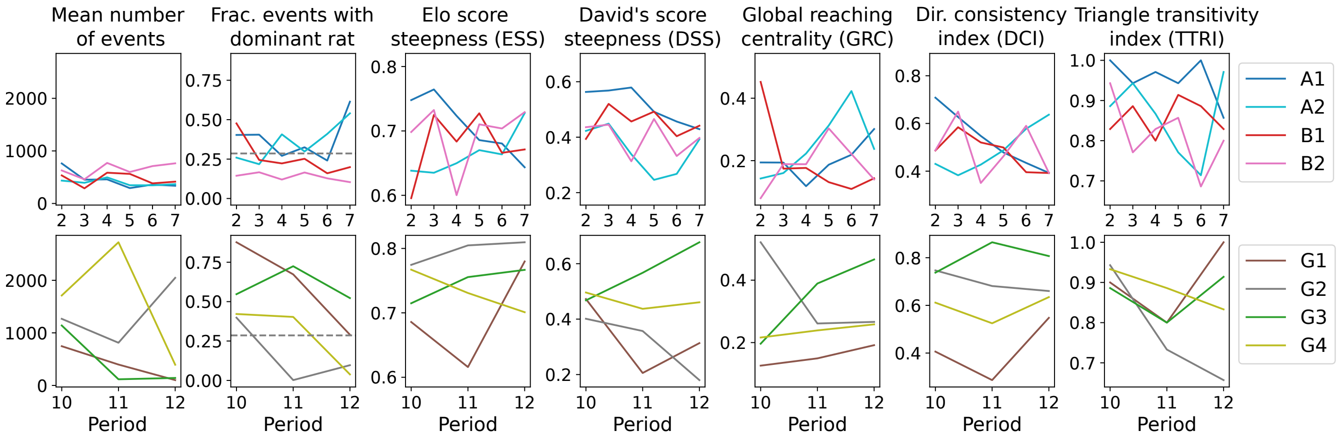


**Figure S7.** **Group social structure metrics over time**. See also Fig 4 for boxplots of the same results, organized by group. The top row shows results for phase 1, and the bottom row for phase 3. The dashed line for fraction of events with dominant rat shows the expected value if all pairs of rats have the same number of events.

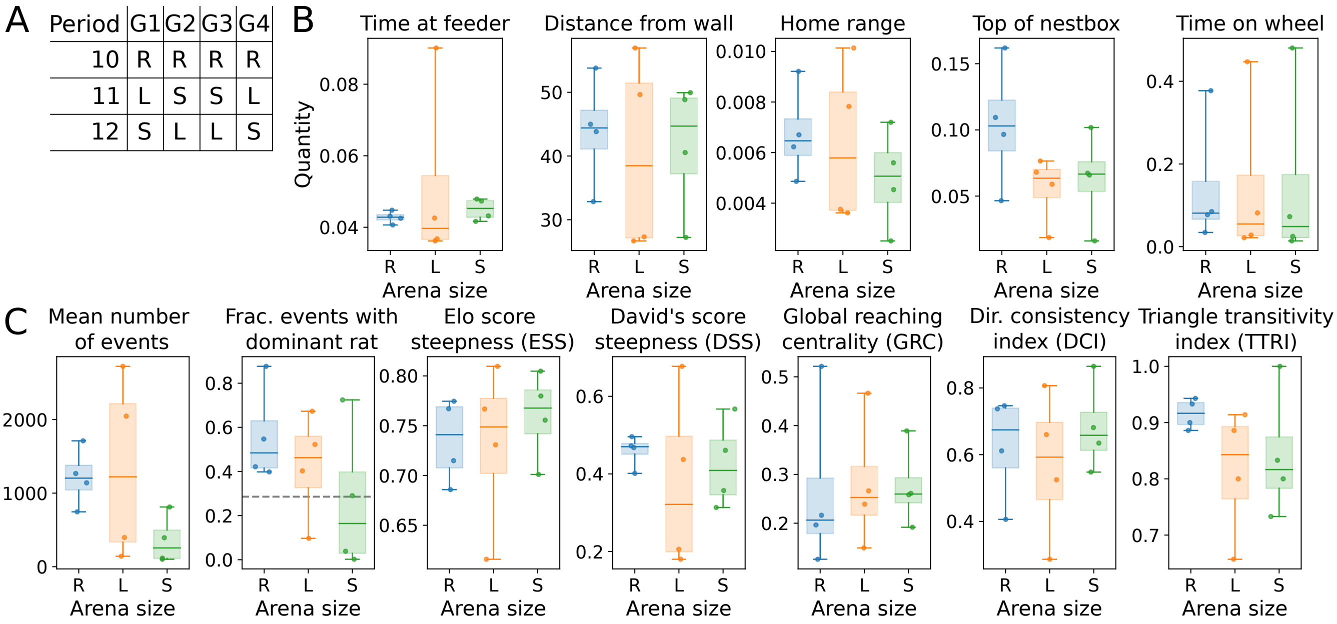


**Figure S8.** **Phase 3 living area size changes, space use, and group social structure metrics**. During phase 3, the size of the living area available to each group was changed by moving the barriers separating the compartments. During Pd 10, all groups had the same Regular (R) - sized living area. In Pds 11 and 12, the living areas were either large (L) or small (S). (A) The living area sizes of each group during phase 3. (B) Space-use metrics according to living area size. (C) Group social structure metrics according to living area size. The dashed line for fraction of events with dominant rat shows the expected value if all pairs of rats have the same number of events.

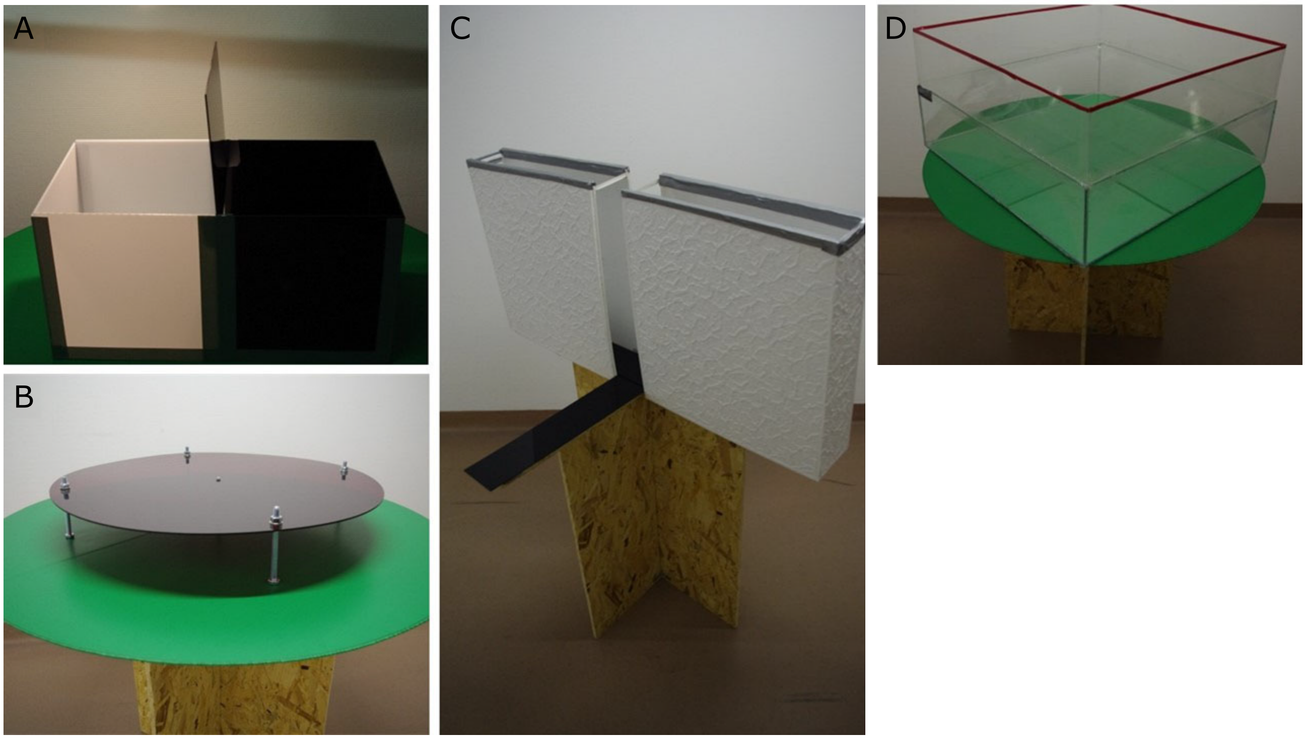


**Figure S9.** **Experimental setup for individual and social tests**. (A) black and white box, (B) canopy, (C) elevated plus-maze, and (D) test apparatus used for pairwise social tests.

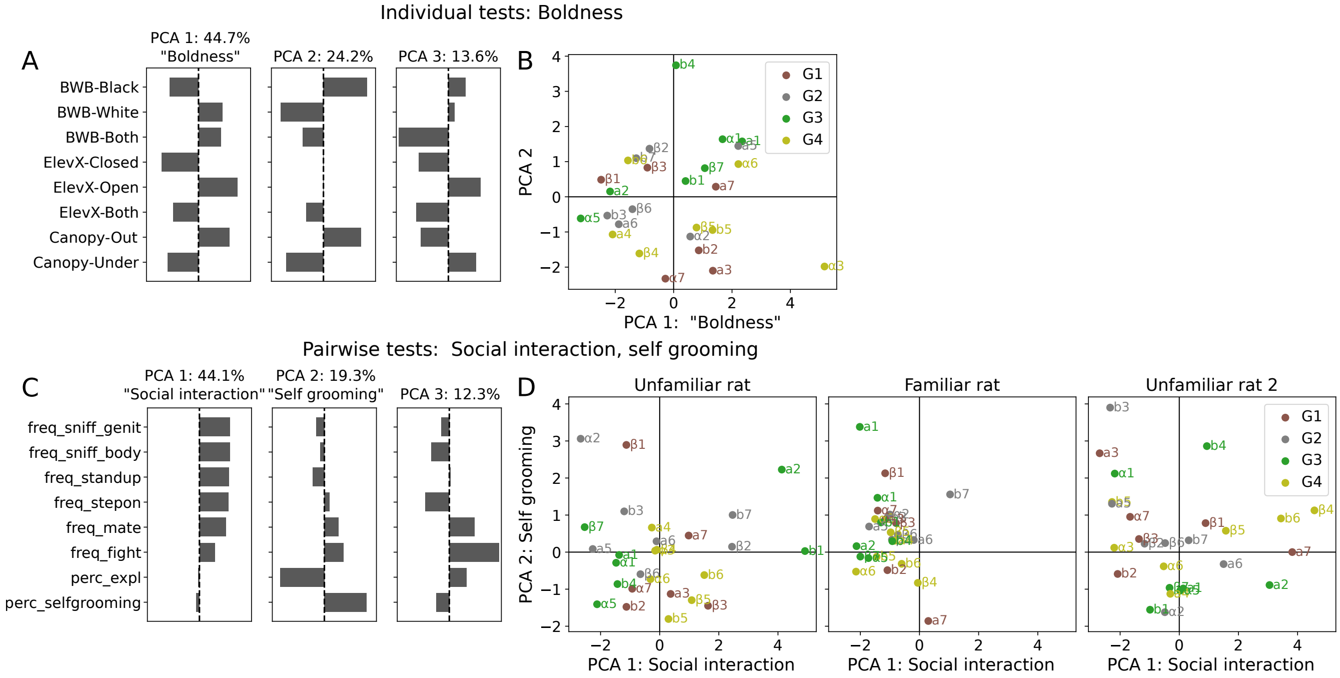


**Figure S10.** **PCA to get individual scores from assays.** (A) The black and white box, canopy, and elevated plus-maze test scores (Figure S9A-C) were used as input to principal component analysis (PCA). The first PCA component is used to define the composite "Boldness" score. (B) Embedding labeling each individual rat's score values projected onto the first two PCA axes, with colors representing the different phase 3 groups. (C) Measures from the pairwise interaction tests with an unfamiliar individual (Figure S9D) were used as input to principal component analysis. Note that while the pairwise tests were also done with a familiar rat, and again with a different unfamiliar rat, the PCA components are determined and set from the first unfamiliar rat tests and these are shown here. Projections onto the first PCA component are used as a "Social interaction" score, and onto the second component as a "Self grooming" score. (D) Embedding labeling each individual rat's scores projected onto the PCA axes shown in (C), for pairwise tests with an unfamiliar individual, a familiar individual, and a second unfamiliar individual. Colors represent the different phase 3 groups.

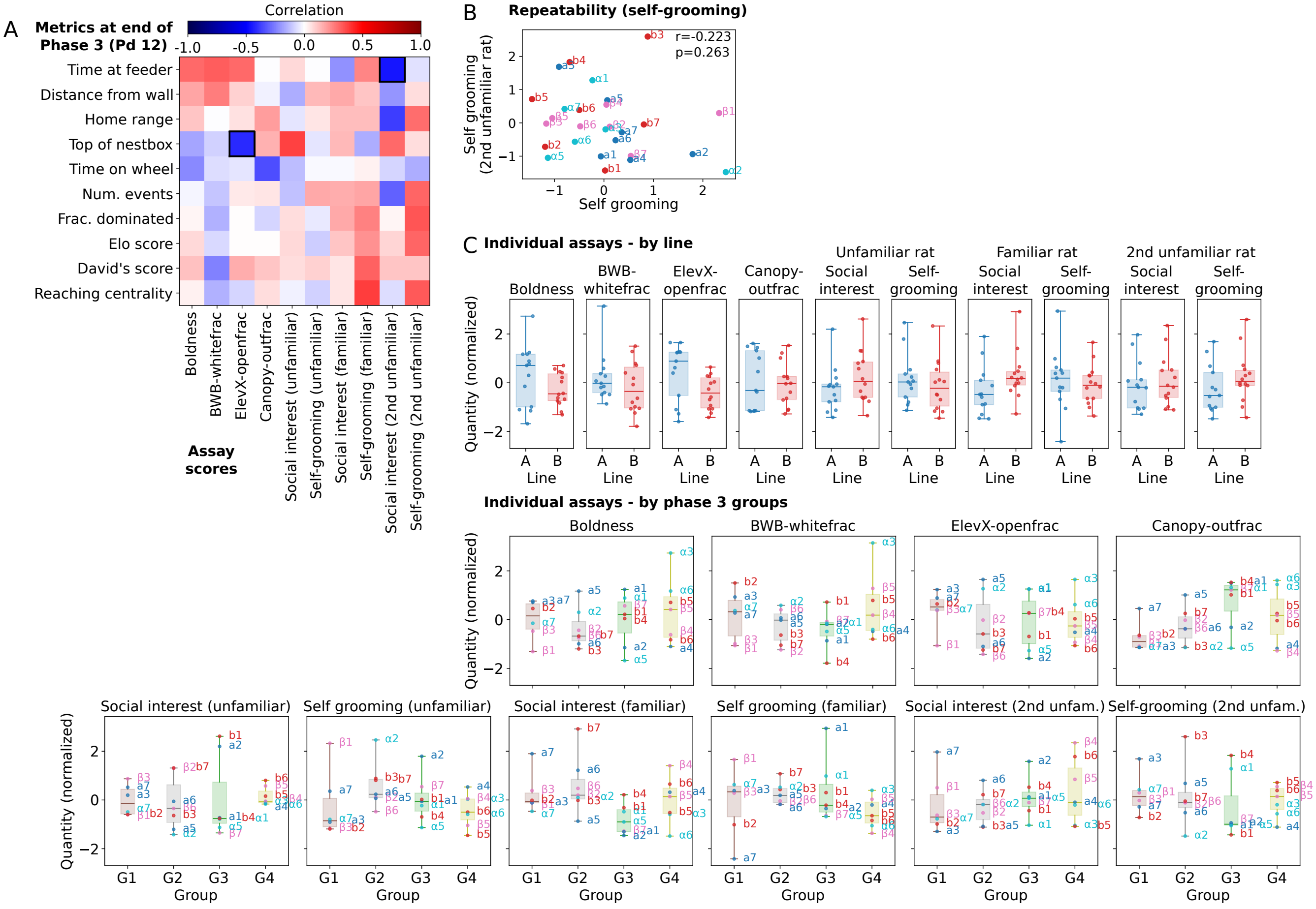


**Figure S11.** **Extended comparison of behavioral metrics to assays**. See Fig 7 for focus comparison with Boldness and Social interaction scores. (A) Pearson correlation values for space use and social behavioral metrics from the final period in phase 3 (Pd 12) with individual assay scores. Labels and color scales denotes correlation values. Statistically significant correlations ($p > 0.05$, calculated using t-distribution) are outlined in bold. (B) Comparison of self-grooming scores (pairwise tests, PCA 2 – see Fig S10C) calculated from tests with a first unfamiliar rat (x-axis), with scores calculated from tests with a second unfamiliar rat (y-axis). See Fig 7 for social interaction score (PCA 1). (C) Individual score distributions according to breeding line (left), and by phase 3 group membership (right). Scores are normalized by the mean and standard deviation of values measured for all rats.

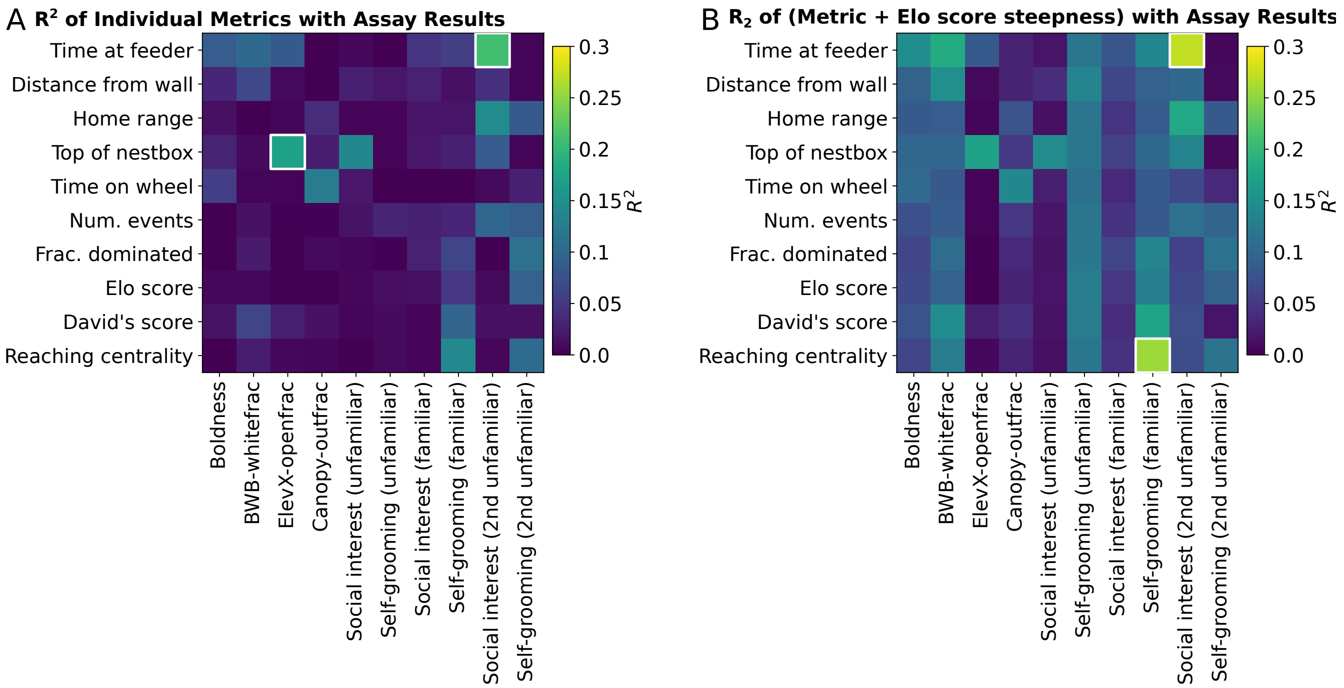


**Figure S12. Interaction of individual and group social structure metrics in predicting individual assay results.** To assess whether interactions between individual and group social structure metrics enhance explanatory power in relation to the behavioral assays, we used linear regression to obtain $R^2$ values that quantify the proportion of variance explained by each model. Each plot shows $R^2$ values, with significant values ($p<0.05$, calculated using an F-test) outlined in white. The model fit results shown are: (A) Each metric fit separately: values displaying the explanatory power of individual space use and social behavioral metrics from phase 3 (Pd 12) for various assay scores. (B) A combined model with each metric and Elo score steepness as the two inputs.

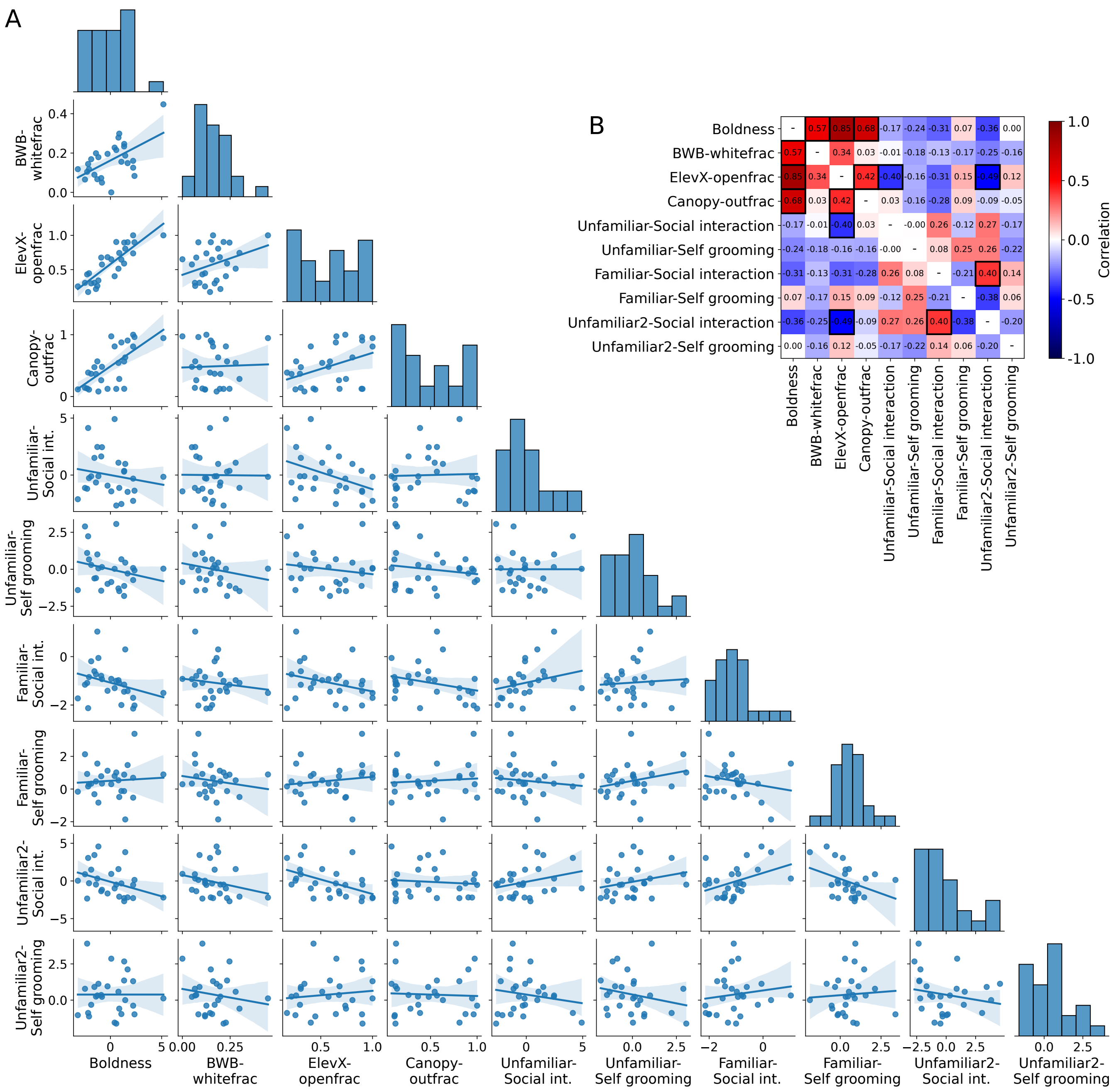


**Figure S13. Comparison of scores derived from individual and pairwise assays.** (A) Scatter plots with regression fit comparing scores derived from the individual and pairwise tests. The boldness score is defined using multiple tests (see Figure S10A), and here is also compared with results from each of these tests. The social interaction and self grooming scores are defined using the pairwise tests (see Figure S10B). (B) Pearson correlation values of scores. The labels and colors indicate correlation values, and bold outlines highlight significant correlations ($p < 0.05$, determined using t-distribution).

**Figure S14. Supplementary video: Time-lapse video of selected days.** Video shows a time-lapse of recorded videos of rats in the experimental arena across 5 experimental days. Preview video online.
